# Theta phase mediates deliberate action switching in human Supplementary Motor Areas

**DOI:** 10.1101/2021.05.06.442965

**Authors:** Giovanni Maffei, Riccardo Zucca, Jordi-Ysard Puigbò, Diogo Santos-Pata, Marco Galli, Adrià Tauste Campo, Rodrigo Rocamora, Paul Verschure

## Abstract

The ability to deliberately overwrite ongoing automatic actions is a necessary feature of adaptive behavior. It has been proposed that the supplementary motor areas (SMAs) operate as a controller that orchestrates the switching between automatic and deliberate processes by inhibiting ongoing behaviors and so facilitating the execution of alternative ones. In addition, previous studies support the involvement of SMAs theta waves (4-9 Hz) in cognitive control. However, the exact role of such oscillatory dynamics and their contribution to the control of action are not fully understood. To investigate the mechanisms by which the SMAs support direct control of deliberate behavior, we recorded intracranial electroencephalography (iEEG) activity in humans performing a motor sequence task. Subjects had to perform a “change of plans” motor task requiring habitual movements to be overwritten at unpredictable moments. We found that SMAs were exclusively active during trials that demand action reprogramming in response to the unexpected cue but were silent during automatic action execution. Importantly, SMAs activity was characterized by a distinct temporal pattern, expressed in a stereotypical phase alignment of theta oscillations. More specifically, single trial motor performance was correlated with the trial contribution to the global inter-trial phase coherence, with higher coherence associated with faster trials. In addition, theta phase modulated the amplitude of gamma oscillations, with higher cross-frequency coupling in faster trials. Our results suggest that within frontal cortical networks, theta oscillations could encode a control signal that promotes the execution of deliberate actions.

## INTRODUCTION

The ability to deliberately interrupt and overwrite ongoing actions as a response to external cues is a necessary feature of cognitive control. For example, when driving a car on a known route, we automatically perform highly trained actions to reach a familiar destination. However, if the habitual route is found blocked, the automatic driving routines are promptly interrupted, and a new motor program is deliberately assembled and selected to follow an alternative path to the goal.

The Supplementary Motor Areas (SMAs), in the Medial Frontal Cortex (MFC), are thought to mediate the switch from automatic to deliberate control when a conflict between current and expected contingencies is detected which requires alterations in ongoing actions (1–5). In particular, it has been suggested that the SMAs orchestrate the balancing of automatic and deliberate processes by inhibiting ongoing motor routines to facilitate the execution of alternative deliberate actions (4,5). Animal studies support this hypothesis, for instance, by showing the involvement of neurons in the primate SMAs in either suppressing or promoting action during “change of plans” paradigms (6). Nevertheless, the functional role of SMAs in humans is still under debate. Imaging studies have reported increased activation of the human pre-SMAs and SMAs during action reprogramming but not during automatic action execution (1). This is consistent with rare lesion studies where subjects suffering from focal SMAs damage are able to stop an action but unable to switch between automatic and deliberate control (7). These results suggest that human SMAs are not generally involved in action inhibition but are specifically supporting switching behavior. However, this interpretation seems at odds with findings from human EEG studies showing significant activation of the medial frontal cortex (MFC) during a behavioral conflict. Here, a positive relationship between the power increase in the theta range (4-8 Hz) and response time during high-conflict trials suggests that this frequency band (often termed ‘frontal theta’) could reflect a generic inhibitory mechanism, possibly acting as a “global brake” on the motor system both pro- and retrospectively (i.e. post-error slowing) (8–10). Hence, the exact role of human SMAs in cognitive control is not clear and pointing to a plurality of possible functions.

A second topic of debate pertains to the mechanism through which the SMAs can control action execution. Indeed, although the power of the theta band over the motor system could reflect an inhibitory mechanism, recent evidence suggests that the phasic period of frontal theta is crucially related to performance in cognitive tasks that involve frontal cortical circuits (11). In particular, theta band phase coherence within and across frontal, motor and parietal regions seems to correlate with a higher accuracy in rule-based decision making (12,13) and memory tasks (14), as well as shorter reaction times during attentional paradigms (15). Despite this growing body of evidence, a direct link between theta oscillatory phase and the executive control of action is still missing (although see (16)). In summary, it remains unclear whether the human SMAs are involved in behavioral switching, whether they have an inhibitory or faciliatory role in action reprogramming and what is the neural mechanism is underlying this putative role. To address these questions, we tested the ability to switch between automatic and deliberate control of action in three (pre-operation) epileptic subjects with iEEG medial-frontal implants in the supplementary motor areas (BA6). Subjects performed a variation of the Serial Reaction Time Task (SRTT) (17), a paradigm that requires the execution of a sequence of repetitive visually-guided key-presses, which becomes progressively automated. In a small subset of trials, the automatic sequence is unpredictably interrupted by the appearance of a cue (switch trials), which required the subjects to halt the ongoing action sequence and press an alternative un-cued key (switch trials).

Consistently with the animal literature (4,6), we find that the SMAs are active during switch trials but not during automatic ones. We also observe that the response time at the single trial level can be predicted from the movement related cortical potential (MRCP) (18). Notably, the ability to detect MRCP peaks at the single trial level allowed us to align the iEEG signals recorded from the SMAs to an endogenous neural event predictive of behavior. This was crucial to minimize intra- and inter-subject variability (19) and to unveil the intrinsic neural dynamics independent of an exogenous (i.e. cue presentation) event-locked analysis (12,20,21). We show that the single-trial theta band phase coherence (22,23), i.e. the contribution of a single trial to the overall phase consistency across trials, were predictive of motor performance at the single trial level, with higher phase coherence correlating with shorter reaction times. Neither amplitude nor power of the MRCP had such predictive power. To further investigate the neurophysiological link between low-frequency oscillations and behavior, we computed the cross-frequency coupling between theta phase and the amplitude of higher frequency activity. This analysis revealed a significant increase of theta-gamma phase-amplitude coupling associated with shorter response latencies suggesting a modulatory effect of theta rhythms on local neuronal activity.

Altogether our results directly support the role of human SMAs in the control of action switching. Moreover, they reveal a possible mechanism of cognitive control based on the entrainment of high-frequency neuronal events by low-frequency oscillations in a cognitive control version of the nested frequency theta-gamma code (24). This interpretation provides further support for the growing body of evidence pointing to the role of phase synchrony and modulation in the theta band as a fundamental operational mode of frontal executive function.

## RESULTS

### Task and behavioral results

In order to explore the neural dynamics underlying behavioral performance in deliberate action switching, three human subjects implanted with intracranial electrodes (iEEG) in the supplementary motor areas (SMAs) (fig. 1-B), performed a variation of the serial reaction time task (17,25) (fig. 1-A). The task was composed of two consecutive blocks for a total of 565 trials. The first block (trial 1-60) required the participants to learn to perform a repetitive sequence of key-presses (N=5) on a touch-screen keyboard by tapping on a visual cue (green, presented for 500 msec or until pressed) until they reached automaticity. Automaticity was defined as the decrease of inter-key-interval (IKI) to an asymptotic value, indicating that the subject effectively internalized the motor sequence and did not rely on visual feedback (17) (fig. 1-C). We observed a significant decrease in IKI between early and late training trials (fig. 1-D) (t-test unpaired between trials 1-20 (N=20) and trials 40-60 (N=20): SBJ1, t=5.30, p<10-^05^; SBJ2, t=5.71, p<10-^05^; SBJ3, t=4.08, p<10-^03^) and its variability (fig. 1-E) (t-test unpaired: SBJ1, t=2.21, p=0.04; SBJ2, t=2.45, p=0.02; SBJ3, t=3.28, p=0.004), confirming that the subjects performed the sequence in an automatic manner (26).

**Figure 1.**
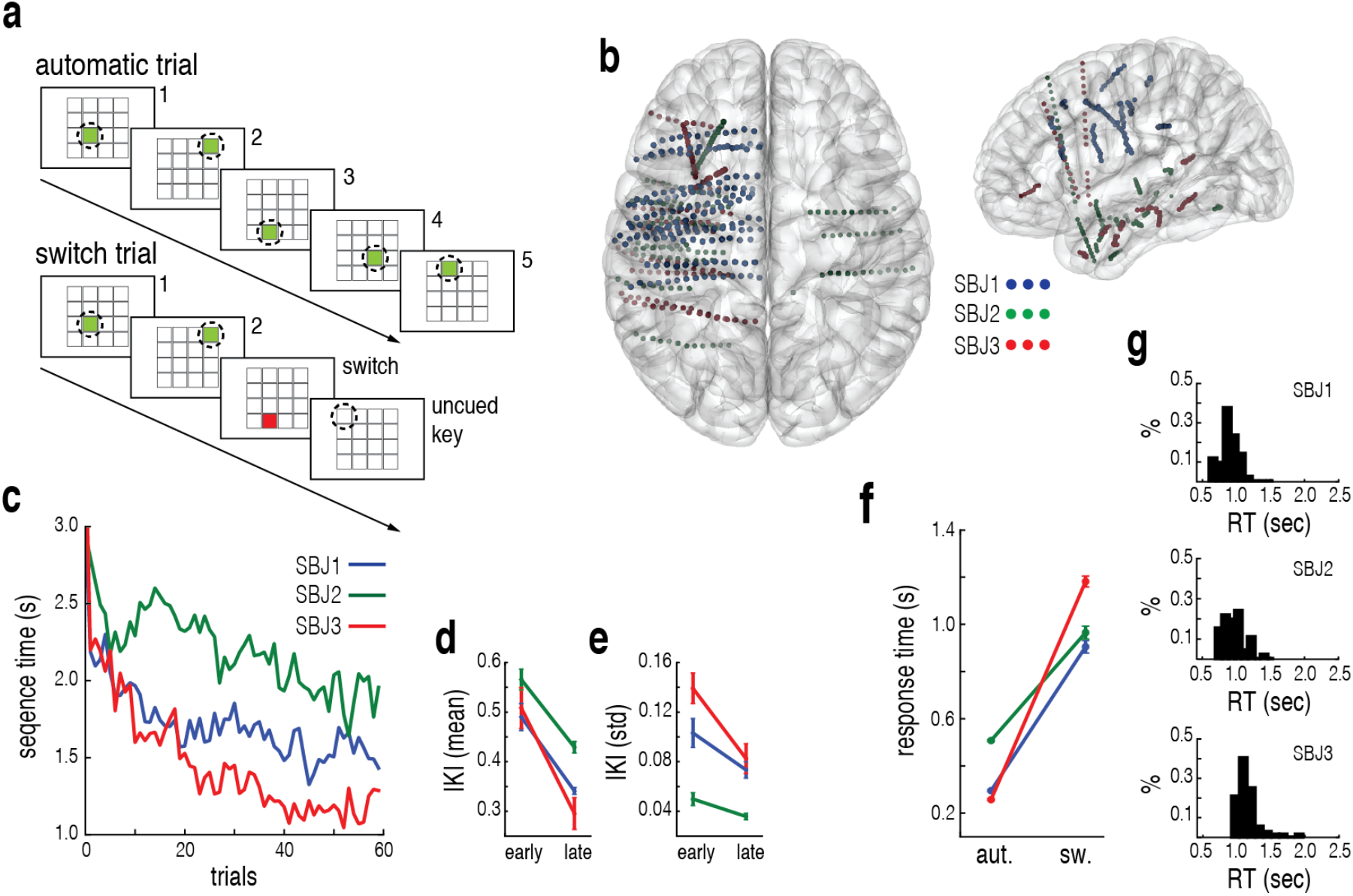
Experimental protocol, recording locations and behavioral performance. **a.** Serial reaction time task. During automatic trials subjects are required to perform a series of visually guided key presses (green key) following a fixed pseudo-randomly generated sequence. During switch trials subjects are presented with a switch cue (red key in a fixed position) appearing at a random step of the sequence, requiring them to interrupt the ongoing motor sequence and press a specific uncued key. **b.** Projection of the locations of relevant contact points in MNI coordinates for each subject. **c.** Evolution of trail duration during training per subject. **d.** Mean Inter-Key-Interval (IKI) in seconds during early (trial 1-20) and late (trial 40-60) phases of training for each subject. **e.** IKI standard deviation during early (trial 1-20) and late (trial 40-60) phases of training for each subject. **f.** Comparison between response time during automatic and switch trials for each subject. Switch responses are collected from all the valid trials in the switch condition from the appearance of the switch cue to the key press. Automatic responses comprise the same number of key presses sampled from the automatic sequence at random from the pool of automatic trials (SBJ 1: N = 43; SBJ 2: N = 46; SBJ 3: N = 72). **g.** Normalized distribution of response time during switch trials for subject. Response time computes as in f.

In the second block of the task (trial 61-565), subjects were required to perform the learned sequence of visually guided key presses. However, these were interrupted by the unpredictable appearance of a switch cue (red circle) at pseudo-random intervals (7 +- 2 sec). When the switch cue appeared, subjects had to interrupt their ongoing automatic motor sequence and press a specific uncued key (fixed for each subject throughout the experiment) as instructed during the training phase (switch trials). Subject 1 and 2 requested to interrupt the experiment prematurely. In total we collected 50 switch trials for SBJ1, 57 for SBJ2 and 80 for SBJ3. Subjects were able to successfully interrupt the ongoing action sequence for most of the switch trials. Switch trials where subjects failed to interrupt the ongoing motor sequence and pressed the next key in the sequence (SBJ1, 14%; SBJ2, 20%; SBJ3, 10%) were excluded from the analysis. The behavioral analysis of the key-presses showed consistently longer response in switch trials compared to automatic response trials (fig. 1-F) (t-test unpaired: SBJ1, t=-39.59, p<0.01; SBJ2, t=-32.96, p<0.01; SBJ3, t=-71.56, p<0.01). This increased latency in response time was accompanied by a greater variability ranging from about 600 to 1300 ms (fig. 1-G). Hence, the motor response dynamics in automatic trials were consistent and stereotyped whereas in switch trials subjects were variably faster or slower than their mean performance at each trial. This difference cannot be explained as a learning effect, as we did not find a significant correlation between trial order and switch response latencies (Pearson’s correlation coefficient: SBJ1: R=-0.11, p=0.18; SBJ2: R=0.08, p=0.37; SBJ3: R=-0.16, p=0.07). In addition, it cannot be consistently explained by the position of the switch cue within the sequence, which showed a significant effect only in one subject (one way ANOVA: SBJ1: F(2,40)=2.546, p=0.091; SBJ2:F(2,43)=2.002, p=0.147; SBJ3:F(2,69)=4.685, p=0.012). This raises the question whether the difference in performance in automatic and switch trials can be explained in terms of the properties of the neuronal process underlying the control of action switching.

### SMAs are involved in switch but not automatic action

It is believed that the SMAs contribute to cognitive control in particular the switching between automatic and deliberate action (2–4). We validated this hypothesis by training a classifier to identify the trial type (automatic vs. switch), by the signals obtained from all the available contact points for each individual subject (see Methods). This classification shows that the signals obtained from the medial-frontal cortex, and in particular the SMAs (BA6C), are most predictive of switch trials (F1 score=0.7) followed by the motor cortex (BA4) (F1 score = 0.57) and the lateral premotor cortex (B6L) (F1 score = 0.52). In particular, we observe that the amplitude of the iEEG local field potentials (LFPs) measured in the SMAs is predictive of the type of trial (automatic or switch) (fig. 2-A, B). Following this step, we restricted our analysis to the relevant contact points in the SMAs in order to determine the precise role of SMAs in switching from automatic to deliberate behaviors.

**Figure 2.**
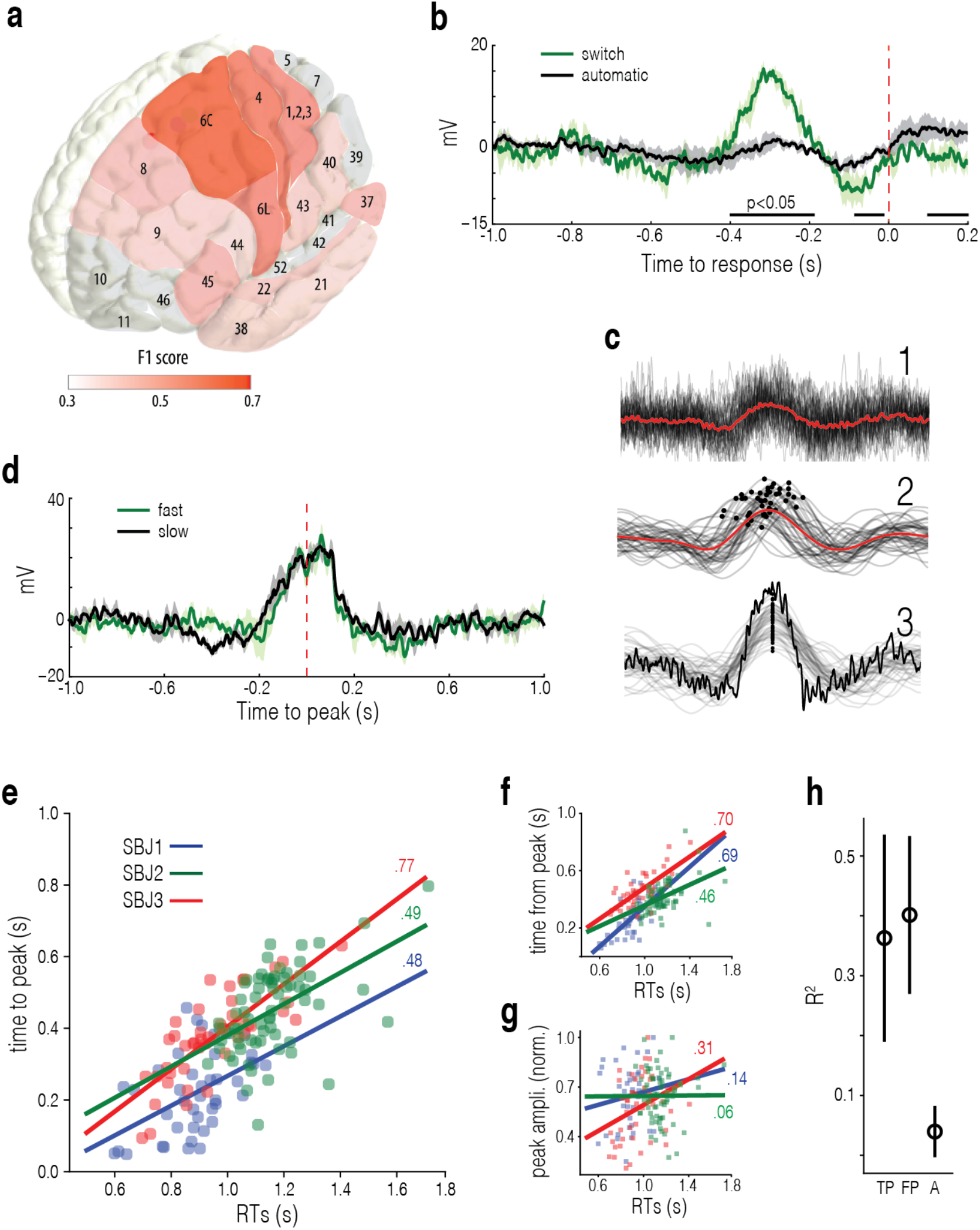
Neural response in SMAs. **a.** Classifier mean prediction accuracy of switch trials projected over a Brodmann atlas. Red hue represents the classification accuracy (F1 score) associated with the areas that were recorded, gray denotes areas not present in the recording. **b.** Motor Related Cortical Potential (MRCP) in SMAs during switch (green) and automatic (black) trials aligned to motor response (red dashed). Switch trials are aligned to the key-press in response to the switch cue (key press of the uncued key), automatic trials are aligned to the last key-press of the automatic sequence. Mean and SEM for each subject. Bar indicates the intersection of significant temporal windows for all the subjects (p<0.05). **c.** Example of alignment of single trial MRCPs to the relative peak for one subject. 1. Key-press aligned MRCPs for each switch trial (black) and mean (red). 2. 2Hz Low passed MRCPs for each trial (black) and mean (red). Black dots indicate the detected peaks. 3. Alignment of low passed filtered MRCPs by their relative peak and unfiltered average. **d.** MRCP in SMAs during fast and slow responses during switch trials (trials sorted by the median of the response time distribution) aligned to the peak of the band-passed MRCP (red dashed). Mean and SEM of subjects. **e.** Relationship between time of response and of cue presentation to MRCP peak during switch trials for each subject. Solid line indicates the linear fit of the data and Pearson’s correlation coefficient. **f.** Relationship between response time and time from MRCP peak to key-press during switch trials for each subject. **g.** Relationship between response time and MRCP peak amplitude during switch trials for each subject. **h.** Mean R^2^ linear regression coefficient between response time and time from cue to peak (TP), time from peak to key-press (FP) and peak amplitude (A). Each score is obtained on the test set as a result of an independent regression for each regressor trained on 70% of the available aggregated trials and tested on the remaining 30%. Mean scores and standard deviation are the result of a 100-fold cross-validation.

Analyzing the time evolution of the SMAs iEEG we observe that during switch trials, a significant movement related cortical potential (MRCP) preceded the response in all three subjects. In contrast, this MRCP was not present during automatic trials (T-statistics cluster permutation (N=1000) analysis: SBJ1, t=34.09, p<10-^4^; SBJ2, t=21.20, p<10-^4^; SBJ3, t=29.69, p<10-^4^) (fig. 2-B). To control whether this MRCP encoded the motor sequence initiation rather than a switch action per se (27), we aligned the LFPs for each subject to the cue indicating the beginning of the automatic motor sequence (not shown). We found no significant increase in amplitude confirming that the earlier observed MRCP was specific to automatic-deliberate switching (T-statistics cluster permutation (N=1000) analysis: SBJ1, t=7.64, p=n.s.; SBJ2, t=4.73, p=n.s.; SBJ3, t=8.13, p=n.s.).

Our results confirm that the SMAs mediate the execution of actions that require cognitive control, such as switch trials and are not involved in the control of over-trained sequential motor responses nor the initiation of automatic sequences (1).

Due to the high resolution of intracranial EEG, we aimed to identify robust MRCPs at the single trial level in order to determine what features of the neural signal were predictive of behavioral performance. To achieve this, we extracted the temporal and amplitude information for each event by band-pass filtering the signal at low frequencies (1-2 Hz) and further extracting the absolute peak (fig. 2-C-1,2) (18,28). We further analyzed the time window from the switch cue presentation to the MRCP peak (TP), the time window from the peak to the key press (FP) and the peak amplitude (A) in relation to the response time at each trial. Temporal analysis of the MRCP showed a marked correlation between its time to peak (TP) and reaction time with respect to cue presentation at the single trial level (Pearson’s correlation coefficient: SBJ1, R=0.481, p<0.001; SBJ2, R=0.779, p<10^-10^, SBJ3, R=0.499, p<10^-5^) (fig. 2-E). A similar positive correlation was found between single trial time-from-peak to key-press (FP) and the response time (Pearson’s correlation coefficient: SBJ1, R=0.698, p<10^-7^; SBJ2, R=0.707, p<10^-8^, SBJ3, R=0.464, p<10^-5^) (fig. 2-F). In contrast, peak amplitude showed no consistent effect on response times and only in one subject reached statistical significance (Pearson’s correlation coefficient: SBJ1, R=0.141, p=0.371; SBJ2, R=0.314, p=0.035, SBJ3, R=0.06, p=0.959) (fig. 2-G). To further quantify the relevance of the time to peak and amplitude aspects of the MRPC for switch behavior we fit a linear model for each variable and computed the explained variance (R^2^) with respect to the response time (fig. 2-H). This analysis confirmed that the temporal dynamics of the MRPC in terms of time-to-peak from cue onset and time-from-peak to response could accurately predict performance while excluding amplitude as a reliable predictor of motor behavior.

We subsequently analyzed whether the profile of the MRPCs showed any behavior dependent modulation by comparing fast and slow switch trials (fig. 2-D). For each subject, trials were aligned to the time of the peak of the band-passed MRCP (as shown in fig. 2-C-3). Further, these trials were sorted into two groups according to the median value of the response time distribution (i.e. fast and slow) in order to obtain an equal number of trials for each group and potential differences between groups tested using cluster-based permutation analysis. This analysis revealed no differences between fast and slow switch trials (T-statistics cluster permutation (N=1000) analysis: SBJ1, t=5.87, p>0.05; SBJ2, t=9.23, p>0.05; SBJ3, t=8.61, p>0.05) suggesting that, although the MRPC is a characteristic neural signature of switch actions, it does not encode information predictive of behavioral parameters such as response time.

Altogether, these results suggest that the SMAs are recruited for deliberate control of switching behavior and that the temporal features of the MRPC, i.e. phase, are decisive in controlling the response time (29).

### Theta phase aligns in faster actions

We hypothesized that oscillatory dynamics in the theta range could constitute a neural signature of cognitive control by mediating deliberate action switching. Previous reports have suggested a strong implication of phase dynamics in cognitive control (11,12,16) suggesting the possibility that stereotypical phase profiles could underlie deliberate action modulation.

To detect potential stereotypical phase patterns underlying the differences in response times (RT), we first aligned the LFPs to the peak of the MRCP. Note that this is not only a necessary step to filter out inter- and intra-subject differences so as to compare phase profiles across conditions (19), but it could also reveal oscillatory patterns that are removed when using exogenous events as reference (8,12,20). Further, we sorted the switch trials for each subject into two classes of equal size (*fast* and *slow*), by splitting the RT distributions by their median value. For each subject and each available contact point, we then calculated ITPC for fast and slow trials and computed the normalized phase coherence difference following the method described in (30) (see Methods).

Class comparison revealed a significantly larger phase alignment in fast trials compared to slow trials localized in the SMAs for all the subjects. To verify the specificity of this result we applied Z-statistics cluster permutation analysis to all the available contact points in order to identify significant differences in ITPC between fast and slow trials (see Methods) and further projected in MNI space the contact points with a significant difference (p < 0.05) in the time window between −0.6 and 0.4 s with respect to the peak of the MRCP (Figure 3-A, see Methods). We observe that, contact points with significantly stronger phase alignment fall in between Brodmann area 6 and 9 for subjects 1 and in Brodmann area 6 for subjects 2 and 3. This effect was found in the theta frequency range between 4 and 8 Hz (fig. 3-C left for an example for each subject) with an onset varying across subjects between 0.5 and 0.1 seconds before the MRCP peak, possibly due to implant location differences (i.e. more frontal for subject 1). Similar theta range variability was detected in the contact points of individual subjects in areas anatomically or functionally related to the SMA. In particular, in the anterior cingulate cortex (ACC) of subject 3 and in the temporal lobe of subject 2 (see Table 1 for a list of the contact points with significant differences (p < 0.05) in phase alignment between fast and slow trials).

**Figure 3.**
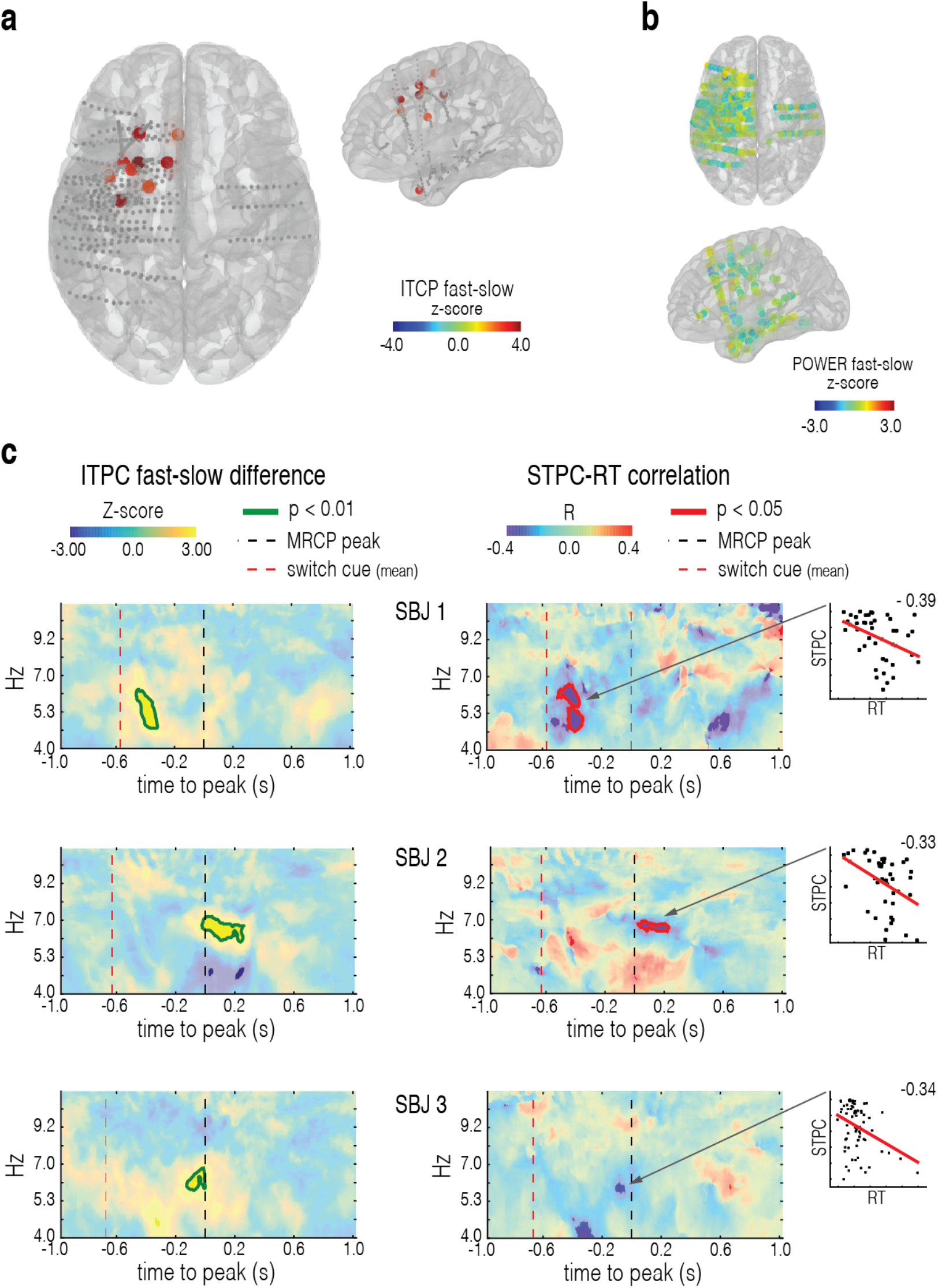
Phase alignment and performance. **a.** Inter-trial phase coherence difference (z-score) between fast and slow (fast-slow) switch trials for the combination of the significant contact points for each subject projected on the MNI space. Contact points showing a significant cluster of time-frequency bins (z > 2.58, Montecarlo p value < 0.05) in the time window −1 s to 1 s centered to the peak of the MRCP are represented in color. **b.** Power difference (z-score) between fast and slow (fast-slow) switch trials for the combination of all the contact points available from all the subjects projected on the MNI space. No contact point reached a statistically significant level, therefore no masking is applied to this figure. **c.** Single subject statistical analysis. **left.** ITPC normalized difference between fast and slow (fast-slow) switch trials. Trials are aligned by the relative peak of the MRCP (black dashed line). Red dashed line indicates the average cue onset time with respect to MRCP peak. Green outline indicates region with Monte Carlo P value < 0.01. **right.** Spearman’s R correlation coefficient between Single Trial Phase Coherence (STPC) and Response Time (RT) computed for each time frequency bin during switch trials. Trials are aligned by the relative peak of the MRCP (black dashed line). Red dashed line indicates the average cue onset time with respect to the MRCP peak. Red profile indicates Monte Carlo P value < 0.05. **box.** Relationship between Response Time and STPC for the significant area shown in the time-frequency plot shown to the left. Red line indicates linear fit.

**Table 1.**
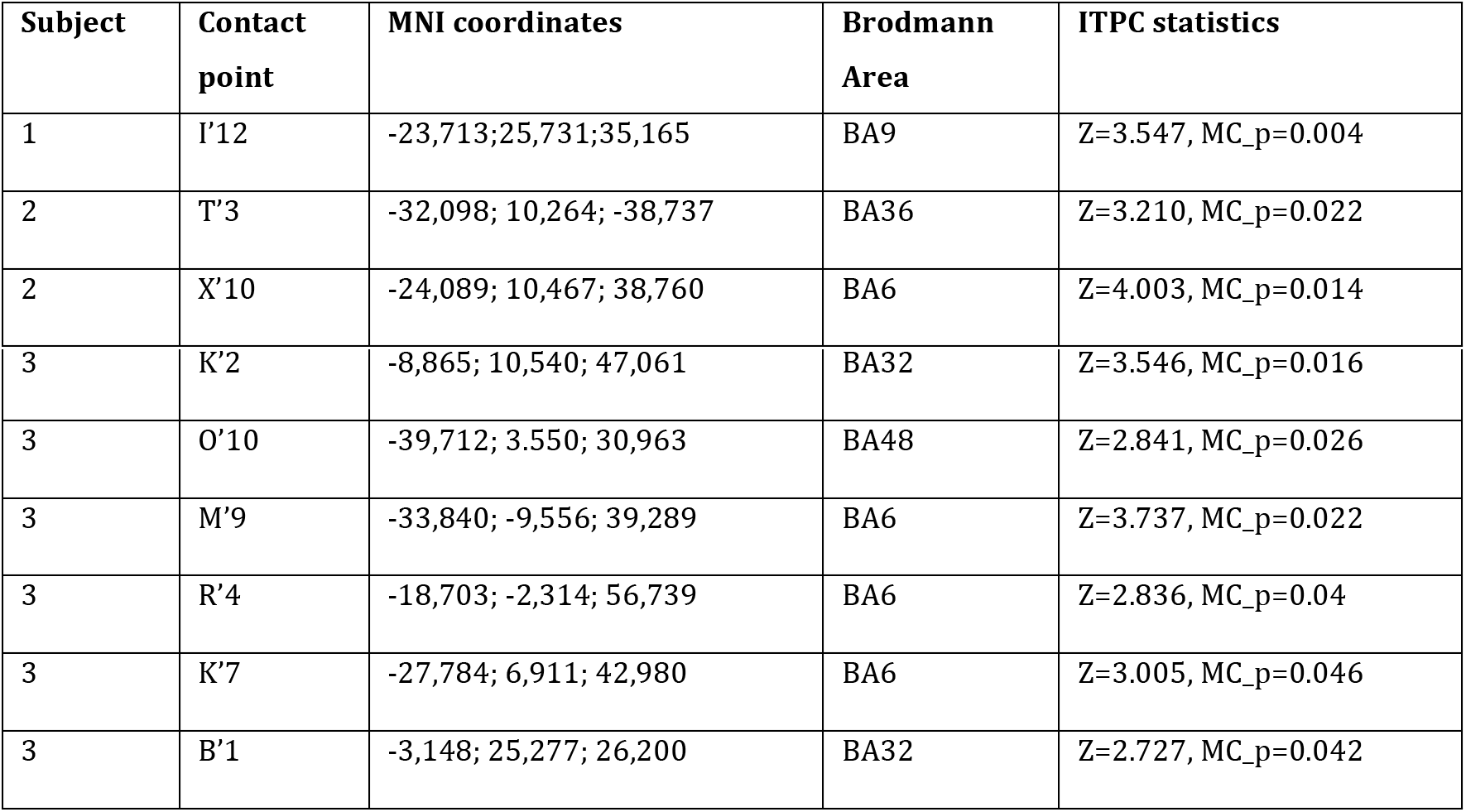
List of all the contact points where the difference in ITCP between fast and slow trials reached statistical significance.

| Subject | Contact point | MNI coordinates | Brodmann Area | ITPC statistics |
| --- | --- | --- | --- | --- |
| 1 | I'12 | -23,713; 25,731; 35,165 | BA9 | Z=3.547, MC_p=0.004 |
| 2 | T'3 | -32,098; 10,264; -38,737 | BA36 | Z=3.210, MC_p=0.022 |
| 2 | X'10 | -24,089; 10,467; 38,760 | BA6 | Z=4.003, MC_p=0.014 |
| 3 | K'2 | -8,865; 10,540; 47,061 | BA32 | Z=3.546, MC_p=0.016 |
| 3 | O'10 | -39,712; 3.550; 30,963 | BA48 | Z=2.841, MC_p=0.026 |
| 3 | M'9 | -33,840; -9,556; 39,289 | BA6 | Z=3.737, MC_p=0.022 |
| 3 | R'4 | -18,703; -2,314; 56,739 | BA6 | Z=2.836, MC_p=0.04 |
| 3 | K'7 | -27,784; 6,911; 42,980 | BA6 | Z=3.005, MC_p=0.046 |
| 3 | B'1 | -3,148; 25,277; 26,200 | BA32 | Z=2.727, MC_p=0.042 |

To further confirm the role of theta phase dynamics with respect to action switching, we applied a single-trial analysis to a subset of contact points in the SMAs that showed the reaction speed and phase alignment effect. Our specific goal was to determine the contribution of inter-trial phase coherence to motor performance. In particular, we computed the total inter-trial phase coherence (ITPC) for each time-frequency bin and inferred the individual contribution of intra-trial coherence by computing a Single Trial Phase Coherence (STPC) pseudo value following (22,23). Finally, for each time-frequency bin, we computed a Spearman’s R correlation between the trial response times (RT) and the relative STPCs (see Methods). We observe a negative correlation between RT and STPC in the 4-8 Hz frequency band once aligned to the peak of the MRCP (fig. 3C, right) (Z-statistics cluster permutation (N=1000) analysis: SBJ1, z>1.96, p<0.001; SBJ2, z>1.96, p<0.001; SBJ3, z>1.96, p=0.07). Altogether these analyses reveal a stereotypic phase profile of theta oscillations locked to the motor potential controlling action switching whose magnitude is monotonically related to the time of the behavioral response at the single trial level.

Importantly, it has been argued that phase coherence may be induced by increases in the power of the oscillations, thus constituting an evoked rather than an actual phase alignment (29). We controlled for this possibility by calculating the difference in power between fast and slow trials and compared the obtained z-scored differences with those obtained in the ITCP domain (fig. 3-B). This analysis confirms a significant effect of ITPC on response time in the absence of significant differences in power suggesting that the detected phase alignment is likely the result of an actual phase coding mechanism within the theta range.

Altogether, these results suggest that the SMAs play a central role in controlling the ability to switch from the execution of an automatic motor sequence to the execution of a deliberate action prompted by an unexpected cue. The enhanced synchronization of theta frequencies in faster switch trials, in absence of power differences, supports the role of phase dynamics as a mediating mechanism that facilitates the execution of deliberate movements. In addition, the differences found in the ACC and temporal lobe support the existence of a functional network involved in switching. Within this network the ACC could be responsible for selectively biasing processes in favor of task relevant information during high conflict switch trials (31). The temporal lobe in turn might encode for arousal or surprise elicited by the unexpected cue (32,33)or support the mnemonic aspects of the decision making process (e.g. retrieving what key should be pressed once the cue is presented) (34).

### Cross-frequency coupling predicts faster movements

We have shown that the speed of deliberate action switching is accompanied by theta phase alignment. However, the direct physiological link needed to support the hypothesis that theta phase dynamics modulate performance via phase-dependent neural activity needs to be established. We sought to answer this question by determining the modulatory effect of theta phase on local high-frequency activity with the hypothesis that higher modulation could support faster switch actions (fig. 4-A), an analysis for which a measure of cross-frequency Phase-Amplitude Coupling (PAC) is particularly suited (35,36).

**Figure 4.**
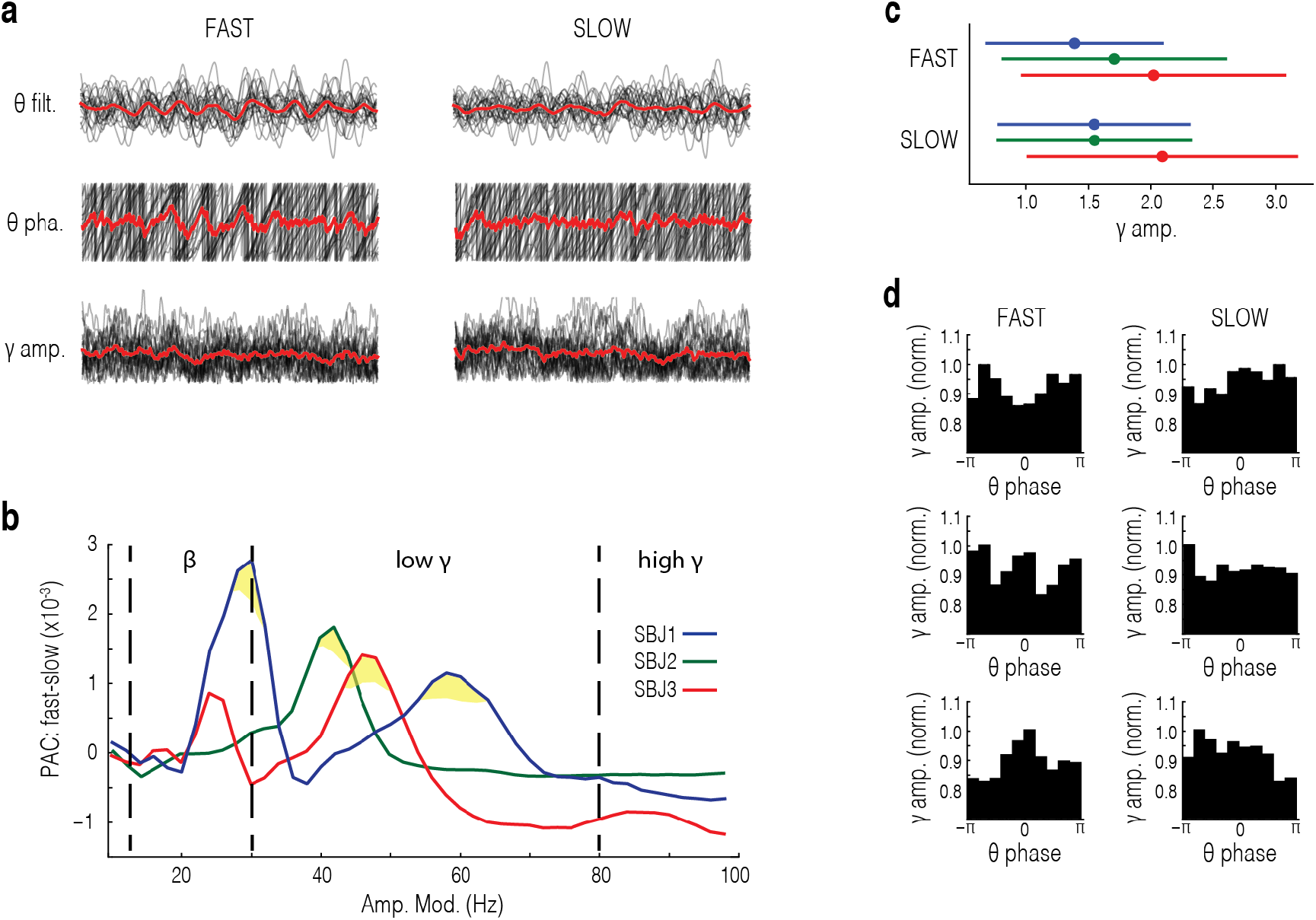
Theta-gamma cross frequency coupling and its relation to performance. **a.** Example of oscillatory activity in SMAs during fast trials (left) and slow trials (right) for SBJ 2. Top. LFP traces of individual switch trials (black) and mean (red) filtered in the theta range (4-8 Hz). Center. Oscillatory phase of the individual LFP traces (black) and mean (red) shown in the above panel. Bottom. Amplitude of the individual LFP traces (black) and mean (red) filtered in the low gamma range (30-80 Hz). All traces are aligned to the peak of the MRCP and sorted by fast (< 900 msec) and slow (> 900 msec) response time. **b.** Difference in phase amplitude coupling (PAC) between fast and slow trials (fast-slow) for each subject comparing the phase of theta oscillations (4-8 Hz) to the amplitude of higher frequency bands (x, 30 Hz, 30-80Hz, and 80-100 Hz). Yellow area indicates *p* < 0.05. **c.** Quantification of mean gamma amplitude in fast and slow trials for each subject selected from statistically significant regions marked in yellow in panel b. Bar indicates standard error of the mean. **d.** Distribution of gamma band amplitude (selected from statistically significant regions in b) over the phase of theta for fast (left) and slow (right) trials for each subject.

We restricted our analysis to the contact points and the temporal windows of approximately 400 ms where a significant increase in phase alignment was detected (Figure 3C). To achieve the temporal resolution necessary for this type of analysis we obtained one surrogate signal for fast and slow trials for each subject by concatenating the respective single trial windows for all the SMAs contact points for each subject (see Table 1 for a list of contact points). Further, we computed PAC values (using the GLM approach (37)) between 4-8 Hz phase (modulatory frequency) and the amplitude of higher frequencies (10-100 Hz in steps of 2 Hz, modulated frequency), and obtained the difference between the two types of trials. Further, to test the significance of the difference between fast and slow trials, we computed the expected difference in PAC under the null hypothesis by shuffling the two types of trials and extracted the z threshold corresponding to p=0.01 (38) (see Methods). We observed an increased modulatory effect of the theta oscillatory phase on the amplitude of frequency bands in the low gamma range (30-80 Hz), which was consistent across subjects (Z-statistics permutation (N=1000) analysis: SBJ1, z>2.58, p=0.027; SBJ2, z>2.58, p=0.034; SBJ3, z>2.58, p=0.029, fig. 4-B,D). In addition, a significant modulatory effect was found in the beta range (i.e. 30 Hz) for one subject (SBJ1, z>2.58, p=0.019). Further analysis revealed no difference in absolute gamma amplitude between trial types (SBJ1, p=0.23; SBJ2, p=0.26; SBJ3, p=0.65), suggesting that the magnitude of phase modulation, rather than activity *per-se*, has a direct effect on controlling performance (fig. 4-D). This result supports a neurophysiological link between theta phase coherence and deliberate control of the action switching through the modulation of high-frequency activity, often interpreted as a correlate of local population activity (35).

## DISCUSSION

Switching from automatic to deliberate behavior in response to environmental cues is a key aspect of adaptive behavior, however, the substrate and dynamics supporting this ability in the human brain are not fully understood. Here, we have addressed this question by analyzing the iEEG of the neocortex subjects engaged in a switch task. We designed a variation of the Serial Reaction Time Task (SRTT) where automaticity is detected as a progressive shift towards faster and more stereotyped movements, indicating that sequences are performed on the basis of an internal habitual representation and allowed to execute autonomously (7,39,40). Using a classifier of the time-domain features of the iEEG we localized a predictive signal of switch trials to the SMAs which confirmed its role in action control. We found that by aligning trials to action initiation, stereotypical phase profiles in the theta range emerged in the SMAs that predicted switch performance at the single trial level. In addition, we showed that theta rhythms modulated high-frequency power in a performance-dependent manner.

In our design, however, we extended the SRTT by introducing a switch cue the required the subjects to interrupt the automated sequence and execute an alternative key-press. This alternative action was un-cued, and it occurred at a low and variable rate, at unexpected positions in the sequence, in order to both avoid habituation and maintain active deliberation. This manipulation is analogous to the switch cue used in (6). However, in our setup, the alternative action was un-cued, to guarantee a minimum amount of deliberation since subjects had to retrieve and implement a specific instruction set from memory (5). During the switch trials, we found significantly longer response times, suggesting a switch from automatic to controlled, deliberate processes (3,6,41). Within the psychological literature, this behavioral marker is known as switch cost (42), interpreted as the result of an “act of control” that involves attentional, mnemonic and motor processes.

We found that activity in the SMAs could successfully predict whether the action was performed deliberately or automatically, supporting the involvement of human SMAs in executive control. This result extends previous evidence from primate studies that demonstrated a lateralized increase in the firing rate of supplementary eye field neurons that exclusively accounted for successful performance during switch but not automatic actions (4). Although the activity profile found in our study is qualitatively similar to the one reported in (6), it remains unclear what the neurophysiological link between low-frequency LFP events and firing rate is. One possibility is that the motor potential we reported (fig. 2-b) is the result of the sum of synchronized afterhyperpolarization events of SMAs neurons contributing to the extracellular field potential (43). This pattern of activity is also consistent with human fMRI studies that reported a consistent activation of the SMAs and the more anterior pre-SMAs during action switching (1,41,44). For example, SMAs are transiently active, together with the cingulate cortex, during the execution of switch actions triggered by response conflict (1,45). Besides, TMS induced inhibition of medial frontal areas impaired the ability to switch between motor responses but not the ability to initiate action per se, an observation consistent with our results showing no significant change of activity during sequence initiation (46). Moreover, contrary to previous reports (27), we excluded the possibility that the neural signature of switching we report could encode sequence production, as no SMAs activity was found during automatic trials. Instead, we demonstrate, to the best of our knowledge, for the first time the movement-cortical potential (MRCP) as the prominent signature of deliberate action switching in the human SMAs using iEEG. MRCPs are low-frequency potentials generated in association with the planning and execution of a cued or self-paced voluntary movement in premotor and motor regions (18). MRCPs can be decomposed in three distinct components: a slow rise component, known as Readiness Potential, which is more prominent in self-paced movements (47,48). A fast decay component following the RP, called negative slope (NS), which peaks at the moment of maximum negativity in concomitance of action initiation (motor potential - MP). Finally, a rebound activity following movement initiation is associated with movement monitoring. In our task, we identified a strong NS component preceding movement initiation but no clear RP, possibly because the switch action was not internally (i.e. voluntarily) generated. Previous literature indeed shows more prominent NS and absence of RP in cue-guided movements compared to self-paced ones (49).

The absence of RP could also support a recent hypothesis by Schurger and colleagues (50). They suggest that this apparent signature of volitional control could be attributed to an artefact arising from the averaging of stochastic fluctuations in the neural activity over trials. Indeed, within the EEG literature MRCPs are typically extracted from averaging across a large number of trials time-locked to action initiation, in order to filter out the noise and account for artefacts introduced by scalp diffusion (51). This procedure, however, generates an artificial build-up in the averaged signal that reflects the integration of noise in a stochastic decision process (SDP) following a drift-diffusion model, where neural activity reaches a “decision threshold” and drives action initiation due to the combination of evidence and stochastic fluctuations. The latter is seen as the source of the variability in response times (52). In our study, the higher resolution and low SNR of iEEG allowed to robustly detect MRCPs at the single trial level directly from the neural tissue and to extract reliable MRCPs features. This allowed for a single-trial analysis precluded in standard EEG analyses usually deployed in the investigation of RP (8,20,21) that support the SDP hypothesis. From a time-domain perspective, the key finding in the single trial analysis of switch trials is that the temporal aspects of the MRCP peak, but not its amplitude, could accurately predict switch response times. This is at odds with previous EEG reports that showed a significant difference in the amplitude of cortical potentials depending on the speed and strength of the movement (53,54). The difference between our and previous results could be due to the different methods employed in the recording. LFPs indeed represent the extracellular activity of neural populations, and they closely track responses in a restricted area of the neural tissue. EEG, in turn, captures a more spatiotemporally extended yet attenuated LFP that integrates several square millimeters of superficial cortical activity and it may reflect more diffuse macroscopic effects (43).

Another methodological advantage of detecting single-trial time domain features is the possibility to align individual trials to the dynamics of endogenous events. This is a necessary step to compare oscillatory patterns across different conditions and subjects, as it has been shown that phase synchrony measures are sensitive to inter-subject and intra-subject temporal variability (19). In addition, previous studies have shown that locking to endogenous events rather than to exogenous ones (i.e. stimulus presentation) can reveal important and otherwise hidden dynamics (12). By aligning individual trials to the MRCP peak, we found that phase synchrony in the theta band across trials predicted switching actions, where higher synchrony is associated with faster trials. This result suggests that action initiation is locked to stereotypical phase profiles in the theta range, suggesting a mechanism by which neurons in the medial frontal cortex synchronize to convey cognitive control signals to downstream motor areas. This also defines a contrast between our results and the SDP hypothesis, by virtue of analyzing signals relative to an internal reference, i.e. MRCP, we reveal a systematic phase code orchestrating action control that will appear stochastic when referenced to external or overt events such as stimuli or responses.

Importantly, we have shown that high synchronicity is prominently and consistently observed in the SMAs but significant differences were also observed in the anterior cingulate cortex and the temporal lobe of individual subjects (fig. 2a). This result suggests that the SMAs do not operate in isolation during action switching, but form part of a broader functional network that involves sensory and mnemonic processes. Indeed it has been suggested that the dorsal part of the ACC could play a major role in conflict resolution by facilitating the selection of task relevant information during high conflict decisions (31,55). The consistency between the activity in the ACC and SMAs we observe (fig. 3-a) supports this hypothesis and the existence of a functional connection between the two areas (55,56). In addition, it has been recently shown that the medial frontal cortex can flexibly recruit the medial temporal lobe during decision-making by means of phase synchrony in the theta band (34). This dynamic functional link between mnemonic and motor processes could explain the higher ITPC in the temporal lobe found in our data (fig. 3-A) and suggests the involvement of this area in supporting the mnemonic aspects related to the task (e.g. to recall what action should be performed when an unexpected cue is presented).

More generally, phase coding has been acknowledged in the human hippocampus and temporal lobe in mnemonic processes. In particular, (57) found that oscillations in the theta band reset their cycle after stimulus presentation, leading to strong patterns of inter-trial phase coherence that correlate with memory encoding. Similar phase resets in the theta range are found in frontal circuits with greater synchrony underlying correct trials (58) and are thought to mediate attentional shifts. The temporal patterns found in our study could, therefore, be the result of a switch cue induced phase reset, encoding action related information, where the magnitude of the reset and its entrainment of local activity drives faster motor responses. The cycle of theta oscillations could indeed carry patterns of information by modulating the amplitude of higher-frequencies in the gamma range (24). Theta-gamma code is believed to represent a fundamental information-processing mode of the brain responsible for sequential item encoding long-term and working memory within memory areas (59). Recently, signatures of the theta-gamma code have been found in frontal cortical circuits during cognitive control tasks. (12) for example, found increased theta-gamma phase-amplitude coupling between prefrontal and motor regions when decisions followed higher order rules. Few reports have linked phase coding to single trial behavioral performance, as in (15) where the strength of phase-amplitude coupling in frontal and parietal regions correlated with reaction times during the allocation of visuospatial attention. Similarly, we report distinct theta-gamma coupling modulated by behavioral performance which suggests that theta range activity reflects a cognitive control signal that entrains the gamma frequency activity of local neuronal populations (35).

Altogether these results contribute to a growing body of literature that identifies frontal theta phase coding as a mechanism promoting cognitive control of behavior (11). Low-frequency oscillations indeed could be a means of information exchange across distant populations within the same functional network (13,35). For example, (16) found fronto-parietal theta synchrony in primate in primate to be predictive of the ability to correctly switch from automatic to deliberate control. However, differently from our analysis, they did not find a relationship with the response time. A similar synchronization pattern was found between the medial frontal cortex and the basal ganglia (60).

Low-frequency oscillatory patterns could also be the result of cortical travelling waves in the theta and alpha bands that have been identified as a mechanism through which information propagates throughout cortical networks (61). Importantly, the spatial and temporal consistency of travelling waves in the prefrontal cortex has been shown to have a facilitatory effect on working memory retrieval, with greater synchrony predicting faster responses (21). Our results suggest a similar effect within supplementary-motor circuits with phase synchrony having a facilitatory effect on deliberate behavior. Contrary to this hypothesis, it has been suggested that medial-frontal theta oscillations may also reflect inhibitory control by mediating action-slowing during situations of conflict and error (9). In particular, human EEG studies testing interference tasks reported increased power in the theta range during high-conflict trials correlating with an increase in response time, both prospectively and retrospectively (i.e. post-error slowing) (8–10). Even though we did not explicitly test for this aspect within our paradigm, we cannot confirm the role of theta as a signature of inhibition for three reasons. Firstly, our analysis did not reveal any distinctive role of theta oscillatory power. In addition, theta oscillatory phase alignment was found to reduce rather than increasing the response time on deliberate switch trials. Finally, in our experiment, the detected pattern of synchronized activity emerged by aligning the LFP of each trial to the peak of an action related event. If theta synchronization represented an inhibitory signal, this event would not be strictly dependent on action execution and it would be expected to rise consistently earlier than the peak of the MRCP (62). This discrepancy might again be due to the differences in resolution and specificity of iEEG versus EEG recordings.

Overall, our results suggest that within frontal cortical networks, theta oscillations encode a control signal that promotes the execution of deliberate actions. In particular, we propose that the power and phase of theta oscillations may reflect different functional roles, where the former locally encodes a general conflict signal and the latter serves as a long-range communication channel facilitating cognitive control (63). This proposal is in agreement with recent accounts of cognitive control that essentially see action selection (i.e. facilitation) and inhibition as two faces of the same coin and therefore postulate complementary neural mechanism underlying these functions (64,65) such as the power and phase of theta band oscillations. This further confirms that phase coding is a fundamental representational format deployed by the brain.

## METHODS

### Data collection

Data were collected from three right-handed subjects with intractable epilepsy, temporarily implanted with intracranial electrodes (iEEG) as a part of a pre-operation procedure to localize the seizure focus. Electrode placement was determined by the surgeons based on the clinical need of each patient.

Data were recorded at the Epilepsy Monitoring Unit of the Hospital del Mar, Barcelona, Spain. All subjects provided the informed consent to participate in the study in accordance with the ethical committee of the Pompeu Fabra University as well as Hospital del Mar. All iEEG recordings were performed using a standard clinical EEG system (XLTEK, subsidiary of Natus Medical) with a 500 Hz sampling rate. A uni- or bilateral implantation was performed using 12 to 16 intracerebral electrodes (Dixi Médical, Besançon, France; diameter: 0.8 mm; 5 to 15 contacts, 2 mm long, 1.5 mm apart) that were stereotactically inserted using robotic guidance (ROSA, Medtech Surgical, Inc).

To identify the anatomical position of the electrode contacts we used the 3D Slicer software(66). With the registration tool, we coregistered (rigid body, 6 degrees of freedom) the post-implantation CT scan to the pre-implantation MRI. We then added the electrode fiducials on a glass model of each patient’s brain obtained with the segmentation tool of the Freesurfer bundle (67). To obtain a single model we coregistered all studies on the MNI152 template provided by the Freesurfer bundle using a semi-automated registration process of 3D Slicer. Briefly, we calculated a linear transform with 12 degrees of freedom by superposing and morphing each patient’s brain MRI onto the MNI brain template, then we used the transform matrix to translate, shift, skew and resize all other studies (CT scan, and unaltered MRI) accordingly. Since the 3D Slicer interface shows the MNI coordinates when hovering the mouse pointer, we could identify structures touched by electrode contacts both by visual inspection and by referring to the aforementioned coordinates.

### Behavioral task

The behavioral task was a variation of the standard Serial Reaction Time Task (SRTT), a type of paradigm that promotes automation of sequential motor behavior (17). Differently from the original task, however, here, in a small subset of trials, the sequential automated action was occasionally interrupted by a cue that required the subjects to switch to a different goal instructed at the beginning of the experiment.

The task comprised a maximum of 500 experimental trials preceded by 60 trials of training. There were two types of trials: automatic and switch. Every trial started with a waiting period of 700 ms +-200 ms during which the screen remained blank. After this, subjects were presented with a virtual 4 by 4 square keyboard.

During automatic trials, a sequence of five keys was highlighted sequentially (green cue) upon button-press. Subjects were instructed to press the cued key as rapidly as possible until the end of the sequence. Each trial terminated at the end of the sequence, and the following one started. The sequence was pseudorandomly generated at the beginning of the experiment to respect a spatial uniform distribution over the keyboard and it was maintained constant throughout the experiment. Switch trials started with the same highlighted key as the automatic trials (green cue), and the next step in the sequence was highlighted upon a button press. Differently from automatic trials, however, one of the intermediate steps of the sequence (i.e. step 2-4 selected at random) highlighted in red (switch cue). Upon presentation of the switch cue, subjects were required to halt the ongoing sequence of movements as fast as possible and press an alternative, uncued key. Participants received all the instructions prior to the beginning of the experiment. Feedback was provided for neither the correct nor incorrect performance. The training phase only comprised automatic trials, whereas the experimental phase included a combination of automatic and switch trials pseudo-randomly interspersed every 7 +-2 trials.

The experimental setup ran on a portable capacitive screen fixed to the hospital overbed table. The tablet included a custom-made Java-based application running the experimental task and logged behavioral performance at 50 Hz whereas task synchronization with the neural recordings was achieved through serial communication with the recording system. Subjects sat in a comfortable position that avoided motor constraints to the arm. After receiving the instructions, subjects underwent a short session that exemplified the task. After this, the experimental session started. Subjects could withdraw at any point during the task.

### Electrophysiology pre-processing

All electrophysiological data were preprocessed in Matlab (EEGLAB toolbox) and subsequently analyzed in Python using custom scripts based on the Numpy, Scipy, SkLearn and MNE libraries.

Data were initially filtered using a two-way zero phase-lag, FIR bandpass filter (2-200 HZ) and an additional notch filter (window = 2Hz) at 50Hz, 100Hz and 150Hz to remove AC current contamination and respective harmonics. Following this step, the signals were individually re-referenced to the average potential of all electrodes for each subject.

After filtering, artifacts derived from strong muscle activity or interference due to contact with electrical devices were identified by visual inspection and respective epochs rejected. To reduce remaining artifacts (i.e. cardiac artifacts, muscle twitches), we applied a combination of Principal Component Analysis (PCA) and independent component analysis (ICA). In brief, we performed PCA on all channels and identified those components which accounted for > 98% of the variance. Such components were subsequently decomposed into the same number of independent components through ICA. At this point, each component time series was visually inspected and components that reflected signal artifacts were rejected. The selection of artifact components was based on a careful inspection of their power spectrum, correlation with other physiological measures (i.e. ECG), and the relation to the temporal structure of the experiment. The rejection was performed by setting the ICA weight associated to the artifact component to 0. The signal was further reconstructed by inverting the ICA operation and the subsequent PCA operation after having renormalized the remaining weights.

### Amplitude analysis

For each subject, the filtered and artifact-free signal was split into epochs according to the trial structure of the task. Each epoch was individually baseline corrected by subtracting the mean amplitude value in a temporal window of 500 ms preceding the beginning of each trial. To identify task-selective channels displaying changes in the amplitude of the signal (i.e. Movement Related Cortical Potentials (MRCP)) we extracted a set of 3 descriptors (absolute mean, variance and integral) and assigned binary labels to each epoch according to the trial type (0=automatic, 1=switch). Further, we applied a classification method based on the Linear Discriminant Analysis (LDA) (68). 100 cross-validation steps were performed to assess performance with Fishers F1 score on class-balanced bootstraps of data samples (80% training, 20% testing). The channels providing the highest classification accuracy were finally selected as the task-related channels. Note that this analysis was naive with regards to the electrode location or the polarity of the event. This step allowed us to narrow down our analysis to those contact points that displayed a task-related change in the amplitude (a detectable difference between conditions) for each subject.

Spectral analysis revealed the presence of MRCPs in the low-frequency range between 1 and 2 Hz (not shown). Trial-by-trial MRCP peaks in the switch condition were therefore identified by low-passing the signal up to 2 Hz using a two-ways zero-phase FIR filter and applying a peak detection algorithm that estimated the time of the absolute peak amplitude in the interval between stimulus presentation (switch-cue) and the response. Single-trial stimulus-peak interval, as well as peak-response interval, were further calculated by subtracting the stimulus presentation time from the peak time and the peak time from the response time respectively.

Finally, the statistical analysis of amplitude differences was performed through a T-statistics one-dimensional non-parametric cluster based permutation test (30) as implemented in the MNE toolbox with cluster significance threshold = 0.05 and number of permutations = 1000.

### Spectral Analysis

Spectral analyses were performed using a DPSS multi-taper method (69,70) as implemented in the MNE toolbox. Trials were aligned to the relative MRCPs peak time, rather than to the behavioral response, in order to avoid artifacts due to the averaging temporally variable signal (19). Changes in the power with respect to the baseline where computed by z-transforming the power spectrum. Statistical differences in the time-frequency power between conditions were calculated through T-statistics two-dimensional cluster based permutation analysis as implemented in the MNE toolbox setting cluster significance threshold = 0.05 and number of permutations = 1000 (30).

### Inter-trial phase coherence (ITPC)

We estimated inter-trial phase coherence to quantify the frequency-dependent synchronization across MRCP peak-aligned trials through Phase Locking Value (PLV) method (71). ITCP is computed as:

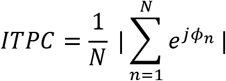

where *N* is the number of trials in one condition and *ϕ* represents the phase estimate at the *n^th^* trial. ITCP is bounded between 0 and 1, where 1 represents full phase synchronization. In order to test differences in ITCP between conditions, we used the cluster-based permutations method proposed by (30). First, we applied a z-transform to the difference in coherence between conditions (Z_ITPC_) that rendered the distribution approximately normal (72):

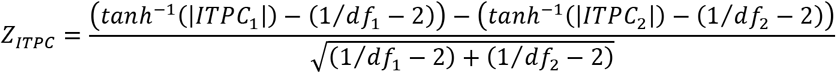

Where *ITPC_i_* and *df_i_* represent the inter-trial phase coherence and degrees of freedom for the *i^th^* condition respectively. To account for the positive bias of ITPC, we used the same amount of trials for the two conditions compared. Second, we selected those regions where z>2.58 corresponding to the 99th percentile of the distribution. Finally, we assessed the significance of the measured difference against the H0 obtained by computing the coherence difference between surrogate groups constructed by permuting 1000 times the original labels and extracting the resulting Montecarlo P value.

### Single trial ITPC

ITPC is by definition an average measure across multiple trials. An estimate of the contribution of the single trial to the average ITPC (STPC), however, can be obtained by computing the difference between the ITPC across all trials and the ITPC across all but one trial following the method proposed by (23) and previously applied by (22). The Single Trial ITPC (*STPC_i_*) for the *i^th^* trial is computed as follows:

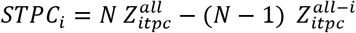

where *N* is the number of trials and 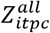 and 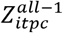 are the z-transformed ITPCs for all trials and all but the *i^th^* trial respectively. Finally, we computed the STPC for each trial and time-frequency bin.

The correlation coefficient between STPC and single trial reaction times was calculated using the Spearman’s R, as the STPC distribution was found to violate the normality assumption. This operation was repeated in order to cover the whole time-frequency range, resulting in an R map of size time - frequency.

To further compute the statistical significance of the obtained R map and to correct for multiple comparisons we applied cluster permutation analysis for each subject. In brief, for each bin we randomly shuffled the data on both the reaction time and the STPC dimensions and recomputed at each permutation the Spearman’s R for a total of 1000 permutations. For each bin we obtained a distribution of R under random condition, which served to set the threshold of significance to the 99% percentile of the distribution, corresponding to a p value of 0.01. Further, each randomly obtained R map was thresholded according to the corresponding significant value, so to obtain a number of time-frequency clusters where the correlation coefficient was found significant. To determine whether the thresholded time-frequency clusters obtained from the experimental condition could be considered statistically significant we compared their magnitude with the ones resulted from the permutation analysis. To this end, for each cluster we integrated the absolute R value so to obtain one magnitude value per cluster. Further we computed the distribution of cluster magnitudes under random condition and calculated its 99% percentile, corresponding to a p value of 0.01. Cluster magnitudes in the experimental condition that exceeded this threshold were considered statistically significant. Whole brain maps were obtained by projecting the average z score of those individual contact points showing a significant cluster in the time window between −0.6 and 0.4 s in the MNI space.

### Phase-amplitude coupling (PAC)

PAC is a measure that quantifies the modulatory effect of low-frequency phase on higher frequency amplitude as a signature of the interaction between their underlying processes resonating at different frequency bands. PAC was computed through the Generalized Linear Models (GLM) method (37) that captures the proportion of variance explained by an underlying linear relationship between analytical amplitude (i.e. envelope, modulated) and phase (modulating) as obtained by Hilbert transforming the signal, using the PACpy toolbox (https://github.com/voytekresearch/pacpy).

We restricted our analysis of PAC to the ROIs emerged from cluster-based permutation analysis and selected as modulatory frequency band the significant frequency domain range for each subject. Our epoch selection was also restricted to the temporal window of approximately 400 ms where a significant increase in phase alignment was detected. For each subject, we obtained one surrogate signal for fast and slow trials by concatenating the respective single trial windows, so to achieve the temporal resolution necessary for this type of analysis. Further, we computed PAC values between the selected modulatory phase and the amplitude of higher frequencies (10-100 Hz in steps of 2 Hz, modulated frequency), and obtained the difference between the two conditions.

Statistical significance between the two conditions was tested through z-statistics against the null-hypothesis of samples from both conditions belonging to the same distribution. This was obtained by randomly permuting the conditions’ labels and calculating the 95 percentile of the maximum PAC value achieved under the assumption that the two conditions were sampled from the same distribution (38). Note that this approach could introduce spurious oscillations as an artifact due to the concatenation of several signals, where the frequency of the oscillation is directly proportional to the length of the segments concatenated. We control for this possibility by choosing temporal windows that may introduce artifacts at lower frequencies than those considered in this analysis. In addition, the same concatenation is applied equally for both conditions and therefore it is unlikely to affect the comparison.

## ACKNOWLEDGMENTS

We thank the patients at the Hospital del Mar - Epilepsy Unit for participating as subjects in the study and the Hospital del Mar - Epilepsy Unit staff for providing valuable technical support during the experiments. This work was supported by the H2020 Research and Innovation grant “Virtual Brain Cloud” (826421). RR and RZ were supported by the project “Clúster Emergent del Cervell Humà” (CECH) ref. 001-P-001682. ATC was supported by the Bial Foundation grant 106/18.

